# Machine learning identifies novel signatures of antifungal drug resistance in *Saccharomycotina* yeasts

**DOI:** 10.1101/2025.05.09.653161

**Authors:** Marie-Claire Harrison, David C. Rinker, Abigail L. LaBella, Dana A. Opulente, John F. Wolters, Xiaofan Zhou, Xing-Xing Shen, Marizeth Groenewald, Chris Todd Hittinger, Antonis Rokas

**Affiliations:** Department of Biological Sciences and Evolutionary Studies Initiative, Vanderbilt University, Nashville, TN 37235, USA; Department of Bioinformatics and Genomics, University of North Carolina at Charlotte, Kannapolis, NC 28081, USA & Center for Computational Intelligence to Predict Health and Environmental Risks (CIPHER), University of North Carolina at Charlotte, Charlotte, North Carolina, USA; Laboratory of Genetics, DOE Great Lakes Bioenergy Research Center, Center for Genomic Science Innovation, J. F. Crow Institute for the Study of Evolution, Wisconsin Energy Institute, University of Wisconsin-Madison, Madison, WI 53726, USA; Department of Biology, Villanova University, Villanova, PA 19085, USA; Guangdong Province Key Laboratory of Microbial Signals and Disease Control, Integrative Microbiology Research Center, South China Agricultural University, Guangzhou 510642, China; Zhejiang Key Laboratory of Biology and Ecological Regulation of Crop Pathogens and Insects, Institute of Insect Sciences, Zhejiang University, Hangzhou 310058, China; Westerdijk Fungal Biodiversity Institute, Utrecht 3584, The Netherlands

**Keywords:** antimicrobial resistance, fungal pathogen, *Candida*, azoles, echinocandins, Erg11, deep mutational scanning

## Abstract

Antifungal drug resistance is a major challenge in fungal infection management. Numerous genomic changes are known to contribute to acquired drug resistance in clinical isolates of specific pathogens, but whether they broadly explain natural resistance across entire lineages is unknown. We leveraged genomic, ecological, and phenotypic trait data from naturally sampled strains from nearly all known species in subphylum *Saccharomycotina* to examine the evolution of resistance to eight antifungal drugs. The phylogenetic distribution of drug resistance varied by drug; fluconazole resistance was widespread, while 5-fluorocytosine resistance was rare, except in *Lipomycetales*. A random forest algorithm trained on genomic data predicted drug-resistant yeasts with 54-75% accuracy. In general, frequency of drug resistance correlated with prediction accuracy, with fluconazole resistance being consistently predicted with the highest accuracy (74.9%). Fluconazole resistance accuracy was similar between models trained on genome-wide variation in the presence and number of InterPro protein annotations across *Saccharomycotina* (74.9% accuracy) and those trained on amino acid sequence alignment data of Erg11, a protein known to be involved in fluconazole resistance (74.3-74.9% accuracy). Interestingly, the top Erg11 residues for predicting fluconazole resistance across *Saccharomycotina* do not overlap with, are not spatially close to, and are less conserved than those previously linked to resistance in clinical isolates of *Candida albicans*. *In silico* deep mutational scanning of the *C. albicans* Erg11 protein revealed that amino acid variants implicated in clinical cases of resistance are almost universally destabilizing while variants in our most informative residues are energetically more neutral, explaining why the latter are much more common than the former in natural populations. Importantly, previous experimental analyses of *C. albicans* Erg11 have shown that amino acid variation in our most informative residues, despite having never been directly implicated in clinical cases, can directly contribute to resistance. Our results suggest that studies of natural resistance in yeast species never encountered in the clinic will yield a fuller understanding of antifungal drug resistance.

## Introduction

Yeasts in the subphylum *Saccharomycotina* (hereafter referred to as yeasts) are genomically diverse, geographically widely distributed, and found in diverse habitats (Opulente et al., 2024). Opportunistic pathogens in this subphylum are a significant global health concern (WHO, 2022), especially for patients with compromised immune systems, for various reasons including for their resistance to antifungal drugs (Lee et al., 2023). For example, initially susceptible strains of *Candida albicans* and *Nakaseomyces glabratus* syn. *Candida glabrata* can quickly evolve (or acquire) resistance to antifungal drugs in clinical settings, whereas other pathogens, most notably the emerging pathogen *Candida auris* are naturally (or natively) resistant (Sanyaolu et al., 2022). Susceptibility screens for antifungal drugs in hundreds of *Saccharomycotina* species have further revealed that a substantial percentage of species are naturally resistant (Desnos-Ollivier et al., 2012). However, even though the genetic variants that underlie evolved resistance in clinical settings have been extensively characterized (Fan et al., 2019; Flowers et al., 2015; Odiba et al., 2022; Wang et al., 2015; Xu et al., 2008), natural genetic variants implicated in antifungal drug resistance are poorly understood.

There are three major classes of antifungal drugs, namely echinocandins (e.g., caspofungin and micafungin), azoles (e.g., fluconazole, voriconazole, and itraconazole), and polyenes (e.g., amphotericin B); as well as two minor classes, allylamines (e.g., terbinafine) and nucleoside analogs (e.g., 5-fluorocytosine), which are used when first line drugs have failed, or in combination with them (Ghannoum & Rice, 1999; Hay, 2023; Lee et al., 2023; Marie & White, 2009; Sigera & Denning, 2023). Resistance to each class has been observed in clinical settings, including observations of pathogens that are resistant to multiple different drugs (Fan et al., 2019; Fisher et al., 2018, 2018; Lee et al., 2023; Marie & White, 2009; Whaley et al., 2016). Elucidating the genetic variants that confer antifungal drug resistance is a crucial step in the development of effective treatment of these pathogens.

One of the main targets of azoles, polyenes and allylamines is the ergosterol synthesis pathway, with resistance in clinical isolates typically conferred through mutations in genes of the pathway. For example, fluconazole resistance in clinical isolates of *C. albicans* is often mediated through mutations in the *ERG11* gene, which encodes a lanosterol 14-alpha-demethylase (Flowers et al., 2015; Odiba et al., 2022). Fluconazole irreversibly binds the active site of Erg11, inhibiting its ability to biosynthesize ergosterol, the primary sterol of the fungal cell membrane. Resistance to azoles can also arise through regulatory changes that may either increase the expression of Erg11 (e.g., via Upc2 (Flowers et al., 2012; Jiang et al., 2016), or that upregulate cellular efflux pathways (e.g., ATP-binding cassette family and the major facilitator superfamily (Marie & White, 2009; Whaley et al., 2016)) to actively remove the drugs. Polyenes also target the ergosterol biosynthesis pathway by binding directly to ergosterol, disrupting membrane stability (Ghannoum & Rice, 1999; Lee et al., 2023). Similarly, terbinafine is an allylamine antifungal that inhibits Erg1 (squalene epoxidase) activity, which also causes a lack of ergosterol and disrupts membrane stability in yeasts (Leber et al., 2003). Resistance to these drugs most often stems from mutations in the drug target(s) that reduce or prevent drug binding (e.g., Erg1) or lead to the production of alternate sterols (Leber et al., 2003, 2003; Lee et al., 2023; Vandeputte et al., 2008). However, these resistance mechanisms often come with severe tradeoffs for membrane stability in yeasts, so resistance to these antifungals is not common (Leber et al., 2003; Lee et al., 2023).

Echinocandins target the cell wall, inhibiting synthesis of β-glucans by binding to the Fks1 and Fks2 proteins, which are important for yeast cell wall resilience (Lee et al., 2023). However, mutations in *FKS1* and *FKS2* that prevent echinocandin binding can reduce drug efficacy (Lee et al., 2023). Finally, 5-fluorocytosine is a nucleoside analog that targets fungal pathogens by inhibiting RNA and DNA synthesis in fungi (Ghannoum & Rice, 1999; Sigera & Denning, 2023). Resistance occurs either by decreased uptake of the drug, or loss of one of the pyrimidine salvage enzymes, which convert 5-fluorocytosine to 5-fluorouridylic acid, the active state of the drug inside the fungal cell (Ghannoum & Rice, 1999; Sigera & Denning, 2023).

Most studies of antifungal drug resistance have been examinations of drug-resistant clinical isolates (Bédard et al., 2024; Flowers et al., 2012; Jacobs et al., 2024; Jiang et al., 2016; Kakeya et al., 2000; Leber et al., 2003; Odiba et al., 2022; Rybak et al., 2021; Wang et al., 2015; Xu et al., 2008). Such studies have been foundational to our understanding of the evolution of antifungal drug resistance and of the genes and pathways involved, but several questions remain. How widespread is natural resistance across entire clades? Can we predict which species are naturally resistant from their genome sequences? And are the genes and mutations that confer natural resistance the same as those that confer resistance in the clinic? To answer these questions, we integrated data from the Y1000+ Project (http://y1000plus.org) encompassing genomic, ecological, and metabolic profiles of over 1,000 yeast species, with experimental measurements of drug resistance against eight clinically relevant, antifungal drugs for hundreds of yeast species and previous experimental and *in silico* deep mutational scanning data of *C. albicans* Erg11 to azole antifungal drugs.

## Results

### The distribution of drug resistance varies across the *Saccharomycotina* yeast phylogeny

To examine patterns of evolution of resistance, we plotted the resistance of 532 yeast species to eight different antifungal drugs (Desnos-Ollivier et al., 2012) on the yeast phylogeny (Figure 1) (Opulente et al., 2024); the majority of strains were natural or environmental isolates (494 of 532 or 93%) with only 38 out of the 532 (7%) isolated from mammalian-associated environments (Table S1). The antifungal resistance profiles of these mammalian-associated yeasts did not significantly differ from the rest of the dataset for any antifungal drug (Table S1). Resistance to fluconazole was by far the most common, with 34.2% (182/532) of species tested being resistant (Table S2). Resistance to voriconazole was the next most frequent (92/532 or 17.2%), followed by caspofungin (69/532 or 13.0%), amphotericin B (53/532 or 9.8%), itraconazole (46/532 or 8.6%), terbinafine (42/532 or 7.9%), 5-fluorocytosine (41/532 or 7.7%), and posaconazole (33/532 or 6.2%) (Table S2). Out of 264 yeasts with resistance to any drug, over half (148) were resistant to two or more drugs. Of the yeasts that were resistant to a single drug, 42.2% (49/116) were resistant to fluconazole (Table S2). We note that each species in our dataset is represented by a single strain. Since differences in resistance between strains of pathogenic yeasts has been observed (Chow et al., 2020; Pais et al., 2022; Selmecki et al., 2006), the resistance phenotype recorded for the strains examined in our study may not be always representative for the entire species.

**Figure 1.**
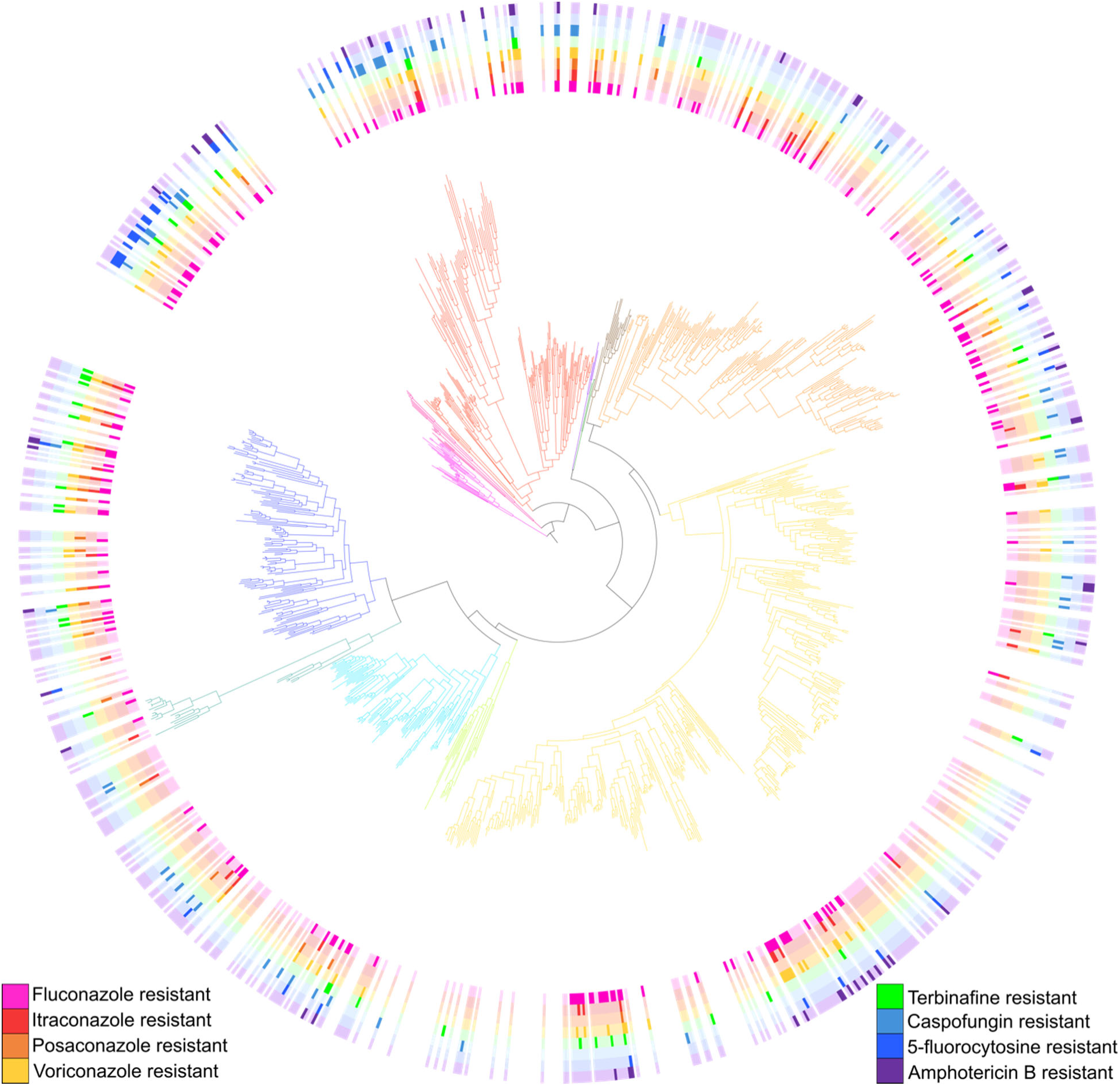
Resistance profiles to antifungal drugs vary throughout the *Saccharomycotina* subphylum. Resistance to some drugs is lineage-specific (e.g., 5-fluorocytosine resistance), but resistance to others is broadly distributed (e.g., fluconazole resistance). Dark colors denote resistance, light colors denote susceptibility, and no color denotes absence of testing. Yeast names are omitted for easier visualization, but they can be found in Figure S2. The colors of the different branches of the phylogeny correspond to the 12 taxonomic orders (Groenewald et al., 2023; Opulente et al., 2024). Drug resistance data obtained using the microdilution technique described in Desnos-Ollivier et al. 2012 (Desnos-Ollivier et al., 2012).

In general, resistance to any of the eight antifungal drugs was rare in the 111 yeasts in the order *Serinales* (which includes the genus *Metschnikowia*, *C. auris*, as well as *C. albicans* and its relatives) (Table S3). Resistance to 5-fluorocytosine was rare outside of the *Lipomycetales* and *Trigonopsidales* orders, while caspofungin resistance was rare only within the *Serinales*, *Pichiales*, and *Saccharomycetales*. In contrast, resistance to the azoles (including fluconazole), terbinafine, and amphotericin B tended to be relatively evenly distributed throughout the phylogeny, with only a few exceptions (Table S3). This pattern of sporadic resistance suggests that resistance to these drugs (or functional analogs of these drugs) was repeatedly gained and lost during yeast evolution. Both the broad distribution and the repeated evolution of antifungal resistance are particularly surprising, considering that 93% of yeast species examined are represented by natural isolates and have never been observed in the clinic (Table S1).

### A random forest algorithm identifies gene and sequence features predictive of resistance

To identify genomic, phenotypic, and ecological features linked to the repeated evolution of drug resistance, we trained a random forest algorithm on genomic, metabolic growth, and isolation environment data from the Y1000+ Project (Harrison, Opulente, et al., 2024; Opulente et al., 2024). Training on metabolic growth and isolation environment data yielded accuracies of 54-75% (average 63%) and 47-63% (average 55%), respectively (Table S4). The features that, on average, most contributed to accuracy for resistance across all the drugs tested were growth on salicin and cellobiose for the models trained on metabolic data, while Arthropoda animal type and having a microbe association were the most informative traits for models trained on isolation environment data (Table S5). The highest accuracy values were obtained when predicting resistance to 5-fluorocytosine in models trained on metabolic data, largely because there are numerous growth substrates (likely unrelated to drug resistance) that show the same clade-specific distribution as 5-fluorocytosine (Table S4). The metabolic and environmental features that most contributed to accuracy for fluconazole resistance were growth on glucosamine and isolation from grasses, respectively (Figure S1).

When trained on genomic data (i.e., on variation in InterPro functional annotations across the genomes of species), models predicted resistance to each of the eight antifungals with ∼54-75% accuracy. Fluconazole resistance was predicted most accurately (74.9%) and itraconazole resistance was predicted least accurately (53.1%) (Figure S2). The higher accuracy in predicting fluconazole resistance likely stems from its high frequency (34%) and broad distribution across *Saccharomycotina* (Figure 1). Anticipating that these features would afford the best potential to uncover insights into the mechanisms of the evolution of drug resistance, we chose to focus on fluconazole resistance going forward.

The most well-characterized genomic determinants of azole resistance in the major human pathogen *C. albicans* are non-synonymous variants in Erg11. Erg11 is the drug target of the azole class of drugs, and resistance can arise by both mutations that impede the drug’s ability to bind to the protein, as well as copy number variants of *ERG11* or its regulators (Fan et al., 2019; Flowers et al., 2015; Kakeya et al., 2000; Kelly, Lamb, & Kelly, 1999; Kelly, Lamb, Loeffler, et al., 1999; Marichal et al., 1999; Odiba et al., 2022, 2022; Sanglard et al., 1998; Wang et al., 2015; A. G. Warrilow et al., 2019; A. G. S. Warrilow et al., 2012). Therefore, we expected to see this gene (or other genes in the ergosterol pathway) in the top features of this model. However, we found that the top features for predicting fluconazole resistance implicated neither Erg11, which ranked 129^th^ in prediction importance, nor other genes in the ergosterol pathway.

Rather, the most informative genomic features were linked to cell wall-associated functional annotations, such as flocculin type 3 repeat (IPR025928; the top feature), which is found in diverse proteins, including in the flocculation proteins Flo5, Flo9, and Flo10 in *S. cerevisiae* (Willaert et al., 2021). These proteins mediate cell-cell adhesion and the formation of multicellular clumps, also called flocs (Willaert et al., 2021). Previous research has found that increasing the number of repeats in these genes linearly increases the adhesion properties of their protein products, as well as the fraction of flocculating cells, which could make cells less accessible to antifungal drugs (Verstrepen et al., 2005). Changes in these genes would also directly affect the composition of the cell wall of yeasts, which could impact the effectiveness of antifungal drugs. Based on the known biology of these genes, we hypothesize that variation in the flocculin type 3 repeats might impact natural drug resistance by changing the structure of colonies of different species and modifying their accessibility to antifungal drugs; although variation in these repeats is unlikely to be a direct mechanism of azole resistance, further testing of this hypothesis would shed light on the relationship between yeast morphological variation and drug resistance.

Variation in the presence/absence in InterPro functional annotations is only one of the many dimensions of genomic variation that differentiate *Saccharomycotina* species. For example, amino acid mutations in the Erg11 protein are the most commonly characterized causes of resistance to fluconazole in clinical isolates of most yeast human pathogens (Flowers et al., 2015; Marichal et al., 1999; Odiba et al., 2022; Rybak et al., 2021; Shahzan et al., 2019; Wang et al., 2015; Xu et al., 2008), yet sequence variation is not accounted for in the InterPro dataset. Therefore, we next focused on predicting fluconazole resistance solely from Erg11 animo acid sequence variation across *Saccharomycotina* yeast species.

### Different random forest models yield similar fluconazole resistance prediction accuracies and implicate the same Erg11 sites

To test whether variation in specific sites of the Erg11 protein sequence contributed to fluconazole resistance prediction accuracy, we identified and aligned Erg11 orthologs across the yeast subphylum using the MAFFT sequence alignment algorithm (Rozewicki et al., 2019). We then trained random forest models to predict fluconazole resistance based on data from (a) both InterPro gene functional annotations and Erg11 MAFFT alignment sites, and (b) just Erg11 MAFFT alignment sites. We found that accuracy of prediction remained similar (75.1% when using both InterPro functional annotations and Erg11 sites, and 73.6% when using just Erg11 sites) (Figure 3). That Erg11 results in similar predictive accuracy as a genome-wide ensemble of functional annotation variation data is consistent with the central role of Erg11 as an azole drug target.

**Figure 2.**
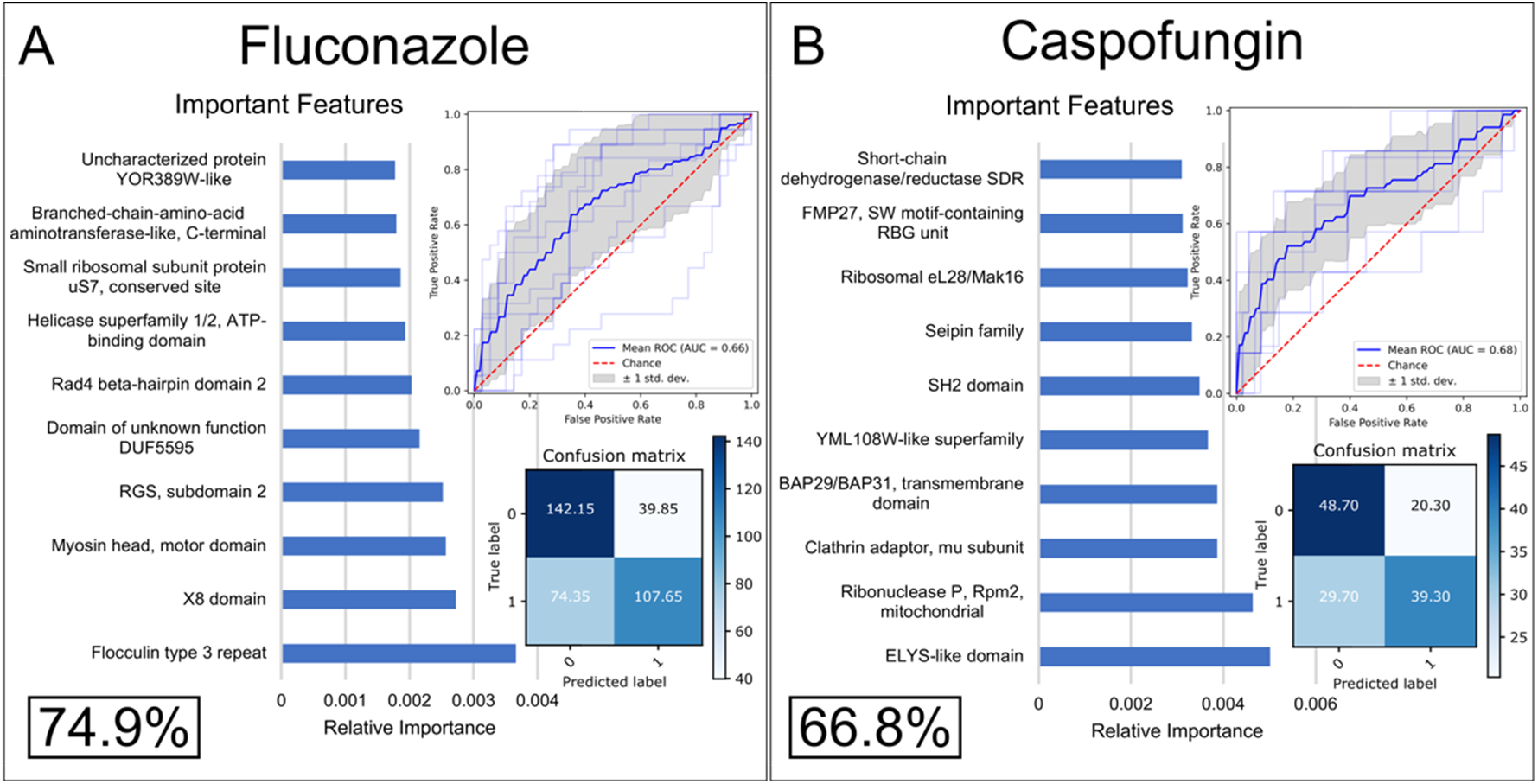
A random forest algorithm predicts resistance to fluconazole (A) and caspofungin (B) with moderate accuracy from variation in InterPro functional annotations. Accuracy is shown in the form of cross-validated balanced accuracy over 20 down-sampled runs (value insight rectangle in bottom left of each panel). Confusion matrices (bottom right) show yeasts predicted correctly to be sensitive (true negatives, top left), yeasts predicted to be resistant but are not (false positives, top right), yeasts correctly predicted to be resistant (true positives, bottom right), and yeasts correctly predicted to be sensitive (false negatives, bottom left). Receiver Operating Characteristic (ROC) curves (upper right) show the true positive rate over false positive rate with changing classification thresholds. Feature importance graphs (left) show the InterPro annotations that are most informative for predicting resistance to each drug. Note that the most informative genomic features were not linked to known drug resistance genes.

**Figure 3.**
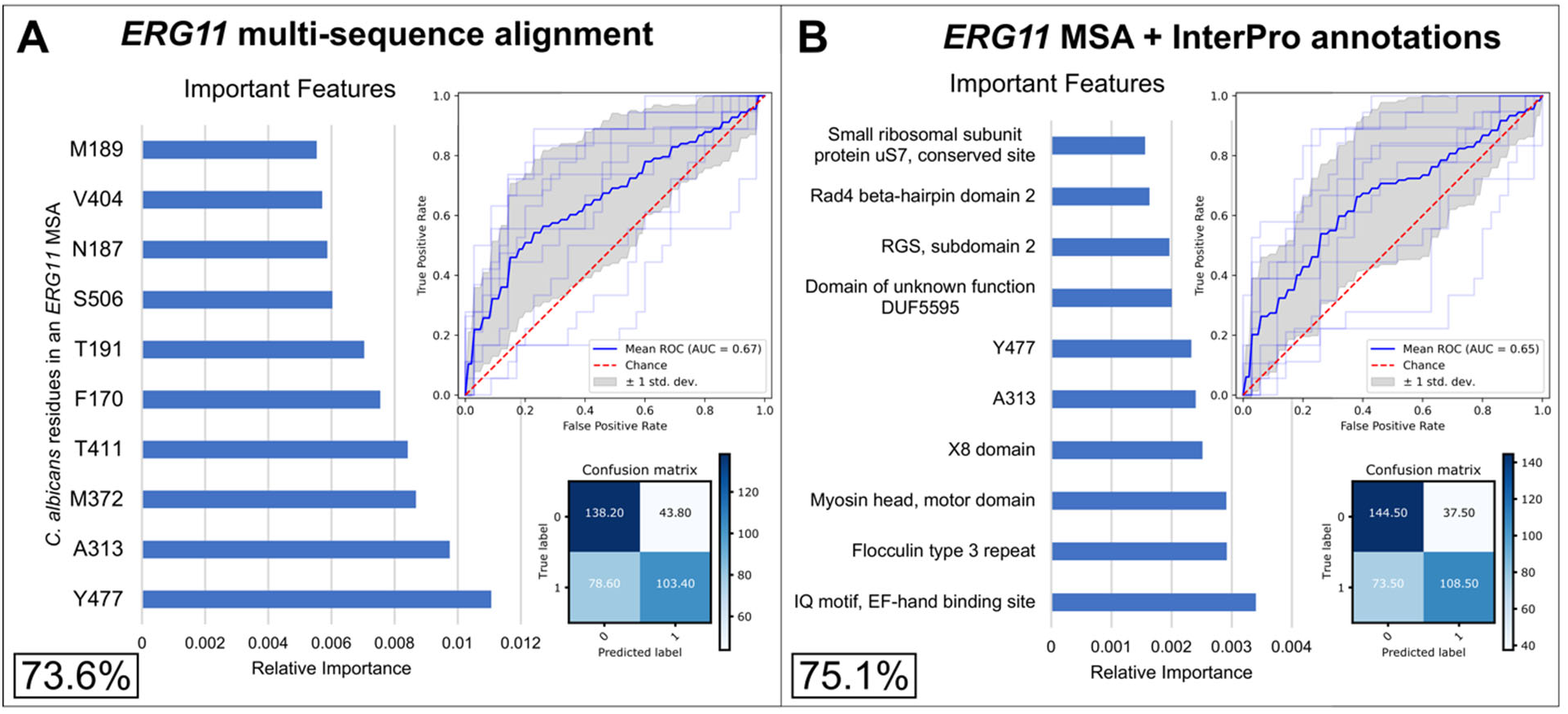
Training a random forest algorithm on the multiple sequence alignment of the known resistance protein Erg11 identifies numerous sites predictive of resistance to fluconazole. Using an integer-encoded multisequence alignment of Erg11 in all *Saccharomycotina* yeasts as input data, the random forest algorithm predicted resistance to fluconazole with moderate accuracy (A). Adding in the InterPro annotations slightly increased accuracy, but residues in the alignment remained some of the most important features (B). Accuracy is shown in the form of confusion matrices (matrix in the bottom right in each panel), which show yeasts predicted correctly to be sensitive (true negatives, top left corner of the matrix), yeasts predicted to be resistant but are not (false positives, top right), yeasts correctly predicted to be resistant (true positives, bottom right), and yeasts correctly predicted to be sensitive (false negatives, bottom left). Receiver Operating Characteristic (ROC) curves (top left in each panel) show the true positive rate over false positive rate with changing classification thresholds. Feature importance graphs (left) show the residues or InterPro annotations that are most informative for predicting growth on fluconazole. The accuracies in the bottom left corner of each panel are cross-validated balanced accuracy over 20 down-sampled runs.

To explore the effects of different methods of aligning and encoding the Erg11 sequence on the training of the random forest algorithm, we used several different methods, including (a) a different sequence alignment algorithm, Muscle5 (Edgar, 2022); (b) one-hot encoding presence and absence of each variant in the alignment; (c) a sequence alignment derived from the superposition of structural models of all Erg11 proteins present in *Saccharomycotina* yeast species; and (d) an alignment-free, k-mer-based (k=3) approach to encode all Erg11 protein sequences from *Saccharomycotina* yeast species. None of these methods substantially influenced prediction accuracy (Figure S3). Importantly, all methods identified many of the same sites in the Erg11 protein as the top predictive features: the three different alignment methods (MAFFT, Muscle5, and structural sequence alignment) all identified same top three most informative sites; and, when using one-hot encoding, seven out of top ten variants were located at 5 sites that were also seen in the top 10 sites of all three alignment methods (Figure S3). Similarly, four of the top five most informative k-mers in the k-mer based method were within two residues of sites in the *C. albicans* Erg11 sequence that were in the top ten most informative sites in the alignment-based methods. These results indicate that diverse methods all identify the same few sites that are most informative for predicting fluconazole resistance.

### Top sites in Erg11 that predict fluconazole resistance experimentally shown to confer resistance across yeasts

To examine whether variation at sites predicted by our models can actually confer fluconazole resistance, we examined data from a recent deep mutational scan experiment measuring the effect of individual amino acid substitutions across 206 sites in *C. albicans* Erg11 on fluconazole resistance (Bédard et al., 2024). Five of our ten most informative sites were tested in these experiments: our top two sites, Y477 and A313, as well as M372, S506, and V404 (numbering based on the *C. albicans* strain CBS 562 protein sequence). Variants at four of those five sites resulted in significantly increased resistance to fluconazole, including variants in the top site, Y477, and in sites A313, S506, and V404 (Figure 4A). Several of these variants are natural variants that differ across *Saccharomycotina* yeasts (e.g., Y477F, A313L, and V404T), demonstrating that natural variants at these sites can confer resistance to fluconazole.

**Figure 4.**
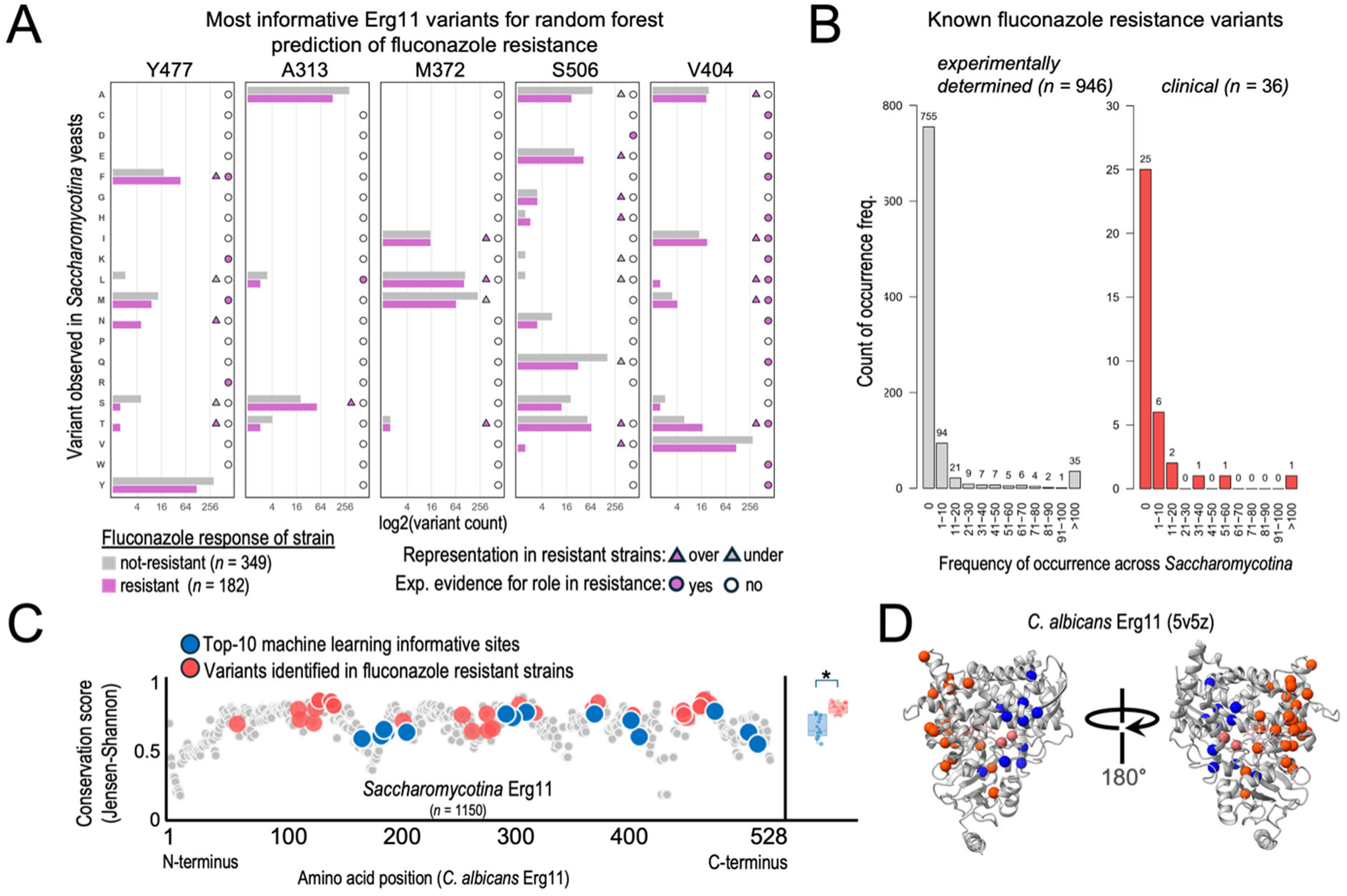
The most informative Erg11 sites are experimentally shown to confer resistance and are generally less conserved than sites previously found to confer fluconazole resistance in clinical isolates. (A) Variant frequencies across Erg11 protein sequences from 1,150 *Saccharomycotina* yeasts for five of the most informative sites identified by a random forest algorithm trained to predict fluconazole resistance. Over- and under-representation of a given variant is shown when either >44% (over) or <24% (under) of fluconazole-resistant yeasts contained the indicated amino acid substitution. Experimental evidence for fluconazole resistance to each amino acid substitution was taken from (Bedard, 2024). (B) Experimentally verified (Bedard 2024) or clinically characterized amino acid substitution frequencies across *Saccharomycotina*. (C) Per-site conservation of all aligned residues of Erg11 across *Saccharomycotina* yeasts. Amino acid positions implicated in fluconazole resistant clinical cases (red) are significantly more conserved than the top ten most informative sites identified by our random forest algorithm (blue) (box plot; p=0.00024, Mann Whitney U Test). (D) Crystal structure of *C. albicans* Erg11 showing spatial distributions of the top ten most informative (blue) and clinical resistance-conferring (red) residues.

There are several other natural variants present at these sites that do not appear to confer fluconazole resistance but differ in their frequencies between drug-resistant and -sensitive yeasts; some of these variants are more common in fluconazole-resistant yeasts, while others are more common in fluconazole-sensitive ones. For example, variant A313S was present in the Erg11 sequences of 71 assayed yeasts and coincided with a fluconazole-resistance phenotype 71.8% of the time; conversely, S506Q was present in 191 yeasts and coincided with a fluconazole-susceptibility phenotype 84.2% of the time. Such patterns of variation, where a variant disproportionality associates with either a resistance or susceptibility phenotype, inform our random forest models and are used by them to predict fluconazole resistance (Figure 4A).

### Top Erg11 sites identified by random forest models are more variable and spatially separated from those conferring fluconazole resistance in the clinic

When using the MAFFT Erg11 multiple sequence alignment to predict fluconazole resistance, the ten sites with the highest feature importance corresponded to *C. albicans* Erg11 residues Y477, A313, M372, T411, F170, T191, S506, N187, V404, and M189 (from most important (0.011 relative importance) to least (0.0055 relative importance)) (Figure 3). Interestingly, none of these residues overlapped with sites harboring 36 Erg11 mutations previously implicated in drug resistance of clinical isolates (Chau et al., 2004; Favre et al., 1999; Flowers et al., 2015; Kelly, Lamb, Loeffler, et al., 1999; Odiba et al., 2022; Rybak et al., 2021; Sanglard et al., 1998; Shahzan et al., 2019; Wang et al., 2015; A. G. Warrilow et al., 2019; A. G. S. Warrilow et al., 2012; Xiang et al., 2013). Twenty-five of the 36 sites had zero importance in predicting fluconazole resistance using the MAFFT alignment, and the remaining 11 had relative importances of 0.0038 or less (Figure 4B, Tables S6 and S7). The minimal contribution of clinically-validated resistance mutations to our predictions reflects the stronger evolutionary conservation at all these sites (mean Jensen-Shannon divergence (JSD) = 0.76) (Table S7). In contrast, evolutionary conservation at our top ten most-informative sites was significantly lower (mean JSD = 0.64; p=0.00024, Mann Whitney U Test; Figure 4C). Only variable sites are expected to be informative for predicting fluconazole resistance in machine learning models, and sites with no variation are simply uninformative for predicting variation in resistance. These results suggest that variants contributing to drug resistance in natural isolates across entire lineages may differ substantially from mutations found to confer resistance in specific pathogens in clinical settings.

Mapping the ten most informative residues for predicting fluconazole onto the high-resolution crystal structure of *C. albicans* Erg11 (5v5z) shows that all ten sites cluster separately from previous clinical variants (Figure 4D). This pattern of spatial segregation, considered in conjunction with the higher sequence conservation of sites that harbor clinical variants across Erg11 protein sequences from *Saccharomycotina* yeasts, suggests that structural constraints may be limiting variation at those sites seen almost exclusively in clinical contexts. Indeed, resistance-conferring clinical mutations all occur within 12Å or less of the Erg11 active site or the natively bound heme (both of which are involved in azole binding), and sites in these functionally important regions are less likely to tolerate variation (mean JSD=0.75 of 213 residues within 12 Å of heme or bound itraconazole in 5v5z). Therefore, differing levels of sequence conservation observed between the ten most informative residues and sites harboring the fluconazole resistance-conferring clinical mutations in *Saccharomycotina* Erg11 proteins may be the result of biophysical constraints in the Erg11 structure itself.

### Erg11 variants informative for predicting fluconazole resistance are less destabilizing than clinical and experimental resistance-conferring variants

Sites harboring resistance-conferring mutations in the clinic are highly conserved and biophysically constrained. In contrast, the ten most informative residues identified by our random forest models are less conserved, raising the question of whether these sites are also biophysically less constrained. To test this hypothesis, we performed an *in silico* deep mutational scan of *C. albicans* Erg11 to evaluate the impact of every possible amino acid substitution on the predicted structural stability of the Erg11 protein (Figure 5A; Methods). We found that Erg11 amino acid variants observed in natural isolates of *Saccharomycotina* are predicted to have significantly lower mean mutational effects per site (i.e., lower changes in their free energy of folding (i.e. ΔΔG)) compared to variants that are never seen (Figure 5B). This distinction also holds when considering these variants individually, with naturally occurring Erg11 variants being substantially less energetically perturbing than variants that are never observed across *Saccharomycotina* (Figure 5C). These observations are consistent with a model of Erg11 protein sequence evolution where purifying selection acts against variants that substantially disrupt the energetic stability of Erg11.

**Figure 5.**
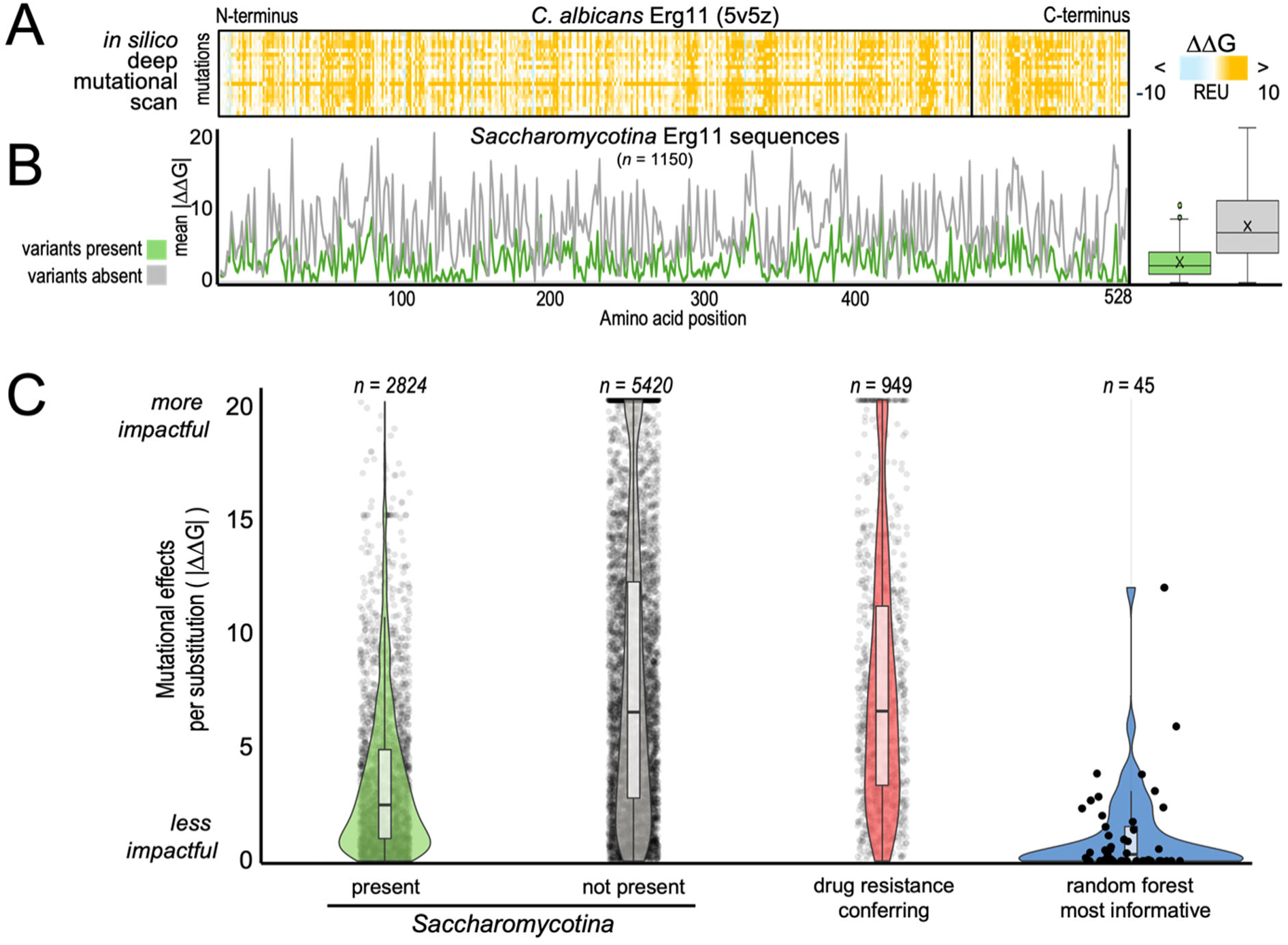
Erg11 variants naturally present in *Saccharomycotina* yeasts, especially those used that best predict fluconazole resistance, are less destabilizing than other Erg11 variants, as well as previously known clinical variants. (A) *In silico* deep mutational scan results of the heme-bound form of *C. albicans* Erg11. Heatmap intensities represent the degree to which each amino acid substitution is predicted to affect the stability of the folded protein relative to wild type (DDG). Positive DDG values are destabilizing, and negative DDG values are hyper-stabilizing. (B) The mean predicted DDG per site, for just those amino acid substitutions that are either present (green) or wholly absent in 1,150 *Saccharomycotina* yeasts. Natural variation is significantly less destabilizing than variation that is never seen (box plot; p=0.00000000, Mann Whitney U Test). (C) Mutational affects for every possible Erg11 mutation. Erg11 variants associated with fluconazole resistance (red) are significantly more destabilizing than those naturally present across *Saccharomycotina*.

Interestingly, if we consider known resistance-conferring mutations identified in the clinic or experimentally determined (i.e., fluconazole resistance-conferring variants identified through an experimental deep mutational scan of 206 sites in *C. albicans* Erg11 (Bédard et al., 2024)), they also are significantly more energetically unfavorable than naturally occurring Erg11 variants present in *Saccharomycotina* yeasts; indeed, they are energetically indistinguishable from those variants that are never observed across *Saccharomycotina*. In contrast to the resistance-conferring clinical and experimental mutations, the predicted mutational effects of the 50 most informative variants from the one-hot encoded model are significantly lower and rank among the most energetically conservative variants seen across *Saccharomycotina* yeasts (Figure 5C).

Thus, while biophysical constraints render clinically-or experimentally-determined fluconazole resistance variants uninformative for predicting resistance from natural Erg11 sequences of *Saccharomycotina* yeasts, machine learning approaches can nevertheless leverage natural variation to accurately predict resistance.

## Discussion

Examining antifungal resistance across 532 *Saccharomycotina* yeasts has informed our understanding of how resistance may evolve outside of clinical settings. Varying levels of resistance to eight different antifungal drugs were observed throughout the subphylum, and a random forest algorithm was effective in leveraging variation in InterPro functional annotation to predict resistance across hundreds of species of yeasts (each represented by a single strain) with moderate accuracy. Genes that impact cell wall composition and colony structure were the most informative, rather than genes known to be directly involved in drug resistance, which suggests that machine learning can pick up on features that impact resistance, even if their molecular mechanism(s) is indirect.

More than one third of *Saccharomycotina* yeasts tested were resistant to fluconazole. This result was a somewhat surprising considering that azoles are synthetic drugs that were developed beginning in the late 1970s (Richardson et al., 1990) and that most of these yeasts were isolated from non-clinical environments. While some of these instances could be due to genomic variation that only incidentally confers azole resistance, azoles have also been widely applied outside of the clinic in agricultural contexts (Fisher et al., 2018). For example, a recent study found that ∼120,000 tons of azoles were sold between 2010 and 2021 just in Europe alone (European Food Safety Authority (EFSA) et al., 2025). Consequently, fluconazole is now routinely found in wastewater, groundwater, surface waters, and drinking water worldwide (Fahy et al., 2025).

Fluconazole directly targets Erg11, and point mutations in Erg11 are known to disrupt drug binding. Many previous studies have identified Erg11 mutations in fluconazole-resistant clinical strains of pathogenic yeasts (Flowers et al., 2015; Odiba et al., 2022; Rybak et al., 2021; Shahzan et al., 2019; Wang et al., 2015; Xu et al., 2008). Therefore, to predict fluconazole resistance, we hypothesized that Erg11 variants in general, and coding variants in particular would be both informative and interpretable. Indeed, an algorithm trained on Erg11 amino acid sequence variation was just as accurate as InterPro functional annotation variation, confirming that variation within Erg11 contributes to prediction of fluconazole resistance. Importantly, previously identified clinical variants were not informative in our machine learning predictions due to their near complete absence across *Saccharomycotina* yeasts. Indeed, sites containing known, azole resistance-conferring residues were among the least variable sites of Erg11. Rather, the most informative residues to our models were among the more variable sites and appear spatially separated from those known sites in the Erg11 protein structure.

To address why that may be, we turned to an *in silico* deep mutational scanning approach to evaluate the impacts of all possible amino acid substitutions to the structural stability of the Erg11 protein. Biophysical modeling of resistance-conferring variants in Erg11 showed that the energetic costs of natural variants observed across *Saccharomycotina* yeasts were much lower than most of the resistance-conferring variants identified in clinical settings or by *in vitro* deep mutational scanning experiments. These results show how machine learning can leverage natural variation at sites proximal to known resistance-conferring sites to predict resistance across large evolutionary timescales.

Our study raises the hypothesis that the variants contributing to natural resistance may be distinct from those that contribute to acquired resistance in the clinic. This idea is supported by the observation that resistance-conferring, single amino acid Erg11 variants observed exclusively within clinical contexts come at large energetic costs, which could reflect strong, short-term selective pressures that are rare or absent in natural populations of *Saccharomycotina* yeasts (Figures 4 and 5). There is extensive support for this hypothesis in studies of drug resistance in bacteria. For example, natural resistance of bacterial species is typically mediated through genetic changes that are distinct from those that confer acquired resistance (Reygaert, 2018).

Furthermore, experimental evolution studies of bacteria grown in the presence of an antibiotic have shown how variation in lifestyle selects for resistance mutations in different pathways; whereas experimental evolution of well-mixed bacterial populations results in the selection of resistance mutations in the protein directly targeted by the antibiotic, evolution of biofilm populations results in the selection of resistance mutations that modulate the regulation of efflux pumps (Santos-Lopez et al., 2019).

The ecological setting for yeast populations evolving drug resistance in natural versus clinical environments is also likely to differ. Drug resistance in the clinic typically evolves because of an infection by a single isolate that propagates inside a patient. This homogeneous pathogen population will likely be exposed to very high concentrations of the drug for long periods of time and throughout a patient’s body (Spagnolo et al., 2021), suggesting that evolving resistance to the drug(s) used to treat the infection is likely to be a main, if not the main, selective agent. In such an environment, a mutation that confers resistance but destabilizes the protein targeted by the drug could be strongly favored. For example, multiple studies in *C. albicans* point to azole-resistance conferring variants being moderately-to-severely compromised in their normal catalytic activity (Kelly, Lamb, Loeffler, et al., 1999; Kudo et al., 2005; Lamb et al., 2000; A. G. Warrilow et al., 2019). In contrast, an environmental yeast is likely to be simultaneously exposed to many more drugs (produced by other microbes), each of which is at a much lower concentration (Chait et al., 2012), as well as to other biotic or abiotic factors. In such a complex environment, large-effect mutations that destabilize protein function would likely be selected against. Rather, natural resistance is more likely to involve mutations of small effect that optimize trade-offs between resistance and protein function.

In 2009, multidrug-resistant isolates of a novel pathogen, *C. auris*, were near-simultaneously identified in multiple continents (Lockhart et al., 2017); *C. auris* has continued its global spread since and is now considered a critical priority fungal pathogen by the World Health Organization (World Health Organization, 2022). The case of *C. auris* emphasizes the importance of understanding the ecology and evolution of lineages harboring fungal pathogens (Rokas, 2022).

If the arguments raised here hold, it follows that the evolutionary pathways to drug resistance are likely to differ between clinical and natural isolates. Thus, large scale analyses of entire lineages can capture natural variation and highlight evolutionary pathways to drug resistance that may be impossible to discover through studies of acquired resistance in the clinic. Such analyses could end up being crucial for identifying drug-resistant species with pathogenic potential *before* they appear in the clinic. We argue that a full understanding of antifungal drug resistance will require examination of both acquired resistance in clinical isolates of yeast pathogens and natural resistance in populations of diverse yeast species that are never encountered in the clinic.

## Methods

### Genomic data matrix

Using InterProScan gene functional annotations generated by the Y1000+ Project (Opulente et al., 2024), a data matrix was built with counts of each unique InterPro ID number in each genome (Table S3). Each genome was its own row, and the number of each InterPro ID (*N =* 12,242) present in one or more of the 1,154 yeast genomes was its own column. A python script recorded the number of each InterPro ID for each genome and put them in the appropriate cells of the data matrix.

### Metabolic data matrix

Our metabolic data matrix contained 122 traits from 893 yeast strains (out of the 1,154 total) from 885 species in the subphylum (Harrison, Ubbelohde, et al., 2024; Opulente et al., 2024) (Table S4). The list of traits in the data matrix included growth on different carbon and nitrogen sources, such as galactose, raffinose, and urea, as well as on environmental conditions, such as growth at different temperatures and salt concentrations. The percentage of missing data in the data matrix was 37.5% (40,906 missing values out of 108,946 total). Less thoroughly studied traits tended to have more missing data than more commonly found and/or thoroughly studied traits.

### Environmental data matrix

The isolation environments for 1,088 (94%) out of the 1,154 yeasts examined were gathered from strain databases, species descriptions, or from *The Yeasts: A Taxonomic Study* (Kurtzman et al., 2011; Opulente et al., 2024) (Table S5) and converted into a hierarchical binary trait matrix using a controlled vocabulary containing all the unique environmental descriptors (Harrison, Opulente, et al., 2024). Strains without isolation environments were either domesticated via crossing or subculturing or lacked information in our searches. The ontology contains six broad isolation environment categories: animal, plant, environmental, fungal, industrial products, and victuals (food or drink). Within these categories, more specific controlled vocabulary annotations are connected to each strain: for example, an isolation environment reported as “*Drosophila hibisci* on *Hibiscus heterophyllus*” is associated in our ontology with the animal subclass “*Drosophila hibisci*” and the plant subclass “*Hibiscus heterophyllus*”.

### Gene sequence data matrix

To retrieve the Erg11 protein sequence(s) from each genome, we used HMMR3 (version 3.1b2) `hmmsearch` (Eddy, 2011). The sequence alignment profile was constructed with `hmmbuild` (Eddy, 2011) from several documented copies of Erg11 in different species across *Saccharomycotina* yeasts. Four of the 1,154 yeasts had annotated copies of Erg11 that were highly divergent and aligned poorly with the others, and were therefore excluded from our subsequent analyses. MAFFT version 7 (Rozewicki et al., 2019) was then used to align the amino acid sequences of Erg11 from the remaining 1,150 yeasts across the subphylum. The resulting multiple sequence alignment was integer-encoded, with each amino acid as well as gaps in the alignment being represented by a different integer, and it was then converted into a data matrix where each column represented a different position in the alignment and each column represented each species. In cases where two copies were found in a genome, the one with the highest sequence similarity score to the HMM profile used to search for Erg11 was used. For the four yeasts with the highly divergent Erg11 sequences, their rows in the dataset were left empty. Muscle5 (Edgar, 2022), as well as a structural alignment (see section on Erg11 structural alignment below), was also used to align the Erg11 protein sequences, and the accuracy, as well as the most important sites in the alignment, remained the same (Figure S3).

### Antifungal resistance data matrix

Our drug resistance data matrix contained eight traits from 532 yeasts in the subphylum. The data were sourced from information available for each of the sequenced strains from the CBS strain database. These data were gathered from strains studied as part of the in the published descriptions of species, additional data on strains obtained by previous studies done in the Westerdijk Fungal Biodiversity Institute (CBS), or additional data provided by the depositors of the strains in the CBS culture collection. The methods for determining whether a strain was resistant are described in Desnos-Oliver et al. 2012; briefly, drug resistance was assessed for each strain using a microdilution technique according to the procedure and criteria established by the Antifungal Susceptibility Testing Subcommittee of EUCAST (AFST-EUCAST) (Desnos-Ollivier et al., 2012).

### Classifying resistance to different antifungals using machine learning algorithms trained on genomic, metabolic, and/or environmental data

To test whether we could classify resistance to eight different antifungal drugs from genomic, metabolic, and isolation environment data, we used a random forest algorithm. For each resistance profile, a random forest algorithm was trained separately on a given dataset to evaluate the accuracy of classification and identify the most important predictive features. Although the task being performed is classification, and the random forest algorithm that we use is a classifier, we refer to the results of this analyses throughout this study as “predictions” for ease of understanding.

We trained a machine learning algorithm built by an XGBoost (1.7.3) (Chen & Guestrin, 2016) random forest classifier (`XGBRFClassifier()`) with the parameters `max_depth=12` and `n_estimators=100`; all other parameters were in their default settings. The `max_depth` parameter specifies the depth of each decision tree, determining how complex the random forest will be to prevent overfitting while maintaining accuracy. The `n_estimators` parameter specifies the number of decision trees in the forest. After testing the increase in accuracy while increasing each of these parameters, we found that having a higher `max_depth` or more decision trees per random forest did not further increase accuracy.

Since drug resistance is typically relatively rare, our datasets tended to be highly unbalanced. Before training the random forest algorithm, down-sampling by randomly choosing an equal number of non-resistant species as resistant species was first employed to balance the datasets. The random forest algorithm was then trained on 90% of the data, and used the remaining 10% for cross-validation, using the `RepeatedStratifiedKFold` and `cross_val_score` functions from the `sklearn.model_selection` (1.2.1) package. Cross validation is a method for assessing accuracy involving 10 trials, each of which holds back a random 10% of the training data for testing. We also used the `cross_val_predict()` function from Sci-Kit Learn separately to generate the confusion matrices; these matrices show the numbers of strains correctly predicted to be resistant or sensitive to a specific antifungal drug (true positives and true negatives, respectively) and incorrectly predicted (false positives, predicted to be resistant but are in reality sensitive; and false negatives, predicted to be sensitive but are in reality resistant). This function also employs a 10-fold cross validation step, but it keeps track of which species are classified as true/false positives and true/false negatives during each of these 10 trials, while entering the final results into a confusion matrix. Top features were automatically generated by the XGBRFClassifier function using Gini importance, which uses node impurity (the amount of variance in resistance for strains that either are or are not resistant to this drug). All these metrics, as well as total balanced accuracy (for which 50% would be equivalent to randomly guessing), were recorded and saved, and then the process was repeated 20 times with new randomly chosen down-sampled datasets each time to account for variation in the yeasts chosen to represent examples of drug-sensitive strains.

Receiver Operating Characteristic (ROC) curves, which plot the true positive rate against the false positive rate, were also generated for each prediction analysis to visualize the accuracy of the algorithm in predicting resistance to a given drug; values of the area under the curve (AUC) greater than 0.5 in these plots indicate better than random classification. Non-down-sampled datasets were used for this analysis, to fully capture the error in the whole dataset.

### Erg11 sequence conservation

The Jenson-Shannon entropy metric of protein sequence conservation was generated from the MAFFT MSA using score_conservation.py (Capra & Singh, 2007).

### Structural alignments of Saccharomycotina Erg11

Hypothetical structural models for all Erg11 proteins found in the Y1000+ Project genomic dataset from the *Saccharomycotina* subphylum were generated using ESMFold as implemented by ColabFold (v.1.5). ESMFold was chosen over other alternative methods (e.g., over methods such as AlphaFold or homology modeling) for its greater speed and comparable accuracy (Lin et al., 2023). A structural MSA was generated from the resulting ESMfold protein models using FoldMason (`foldmason easy-msa --report-mode 1 --refine-iters 5`) (Gilchrist et al., 2024).

### *C. albicans* Erg11 protein structure

A protein structure for the apo form of *C. albicans* Erg11 was retrieved from the PDB (5v5z) (Keniya et al., 2018). The amino acid sequence of the structure was checked and edited to match the Erg11 sequence of the *C. albicans* strain CBS 632 present in the Y1000+ Project (only one amino acid difference was amended). All protein model images were generated using ChimeraX (v1.7) (Meng et al., 2023).

### *C. albicans* Erg11 *in silico* deep mutational scanning

The apo form of *C. albicans* Erg11 (5v5z) was relaxed in complex with the native heme using Rosetta 3.13. Briefly the structure was cleaned and renumbered using `clean_pdb.py` and `pdb_renumber.py`. The cleaned structure was minimized with heme `(-nstruct 20, - relax:cartesian true, -default_max_cycles 200`), and the lowest energy structure was chosen.

All-way, *in silico* mutagenesis (deep mutational scanning) was conducted using Rosetta 3.13 (`cartesian_ddg`) and energy minimization protocols and parameterizations previously benchmarked to optimize replication of experimental ΔΔG (DDG) measurements (`parser:protocol cartesianrelaxprep.xml`) (Frenz et al., 2020). Three replicates were performed for each substitution (`ddg::iterations 3`) and the mean change of free energy of folding (ΔG) was derived from the mean difference between wild type and each amino acid substitution (ΔΔG = ΔG (mutant) - ΔG (wild type)) across replicates.

## Acknowledgements

The authors thank members of the Rokas Lab and Y1000+ Project (http://y1000plus.org) team members for helpful discussions. This project was supported by the National Science Foundation under Grants No. DEB-2110403 (C.T.H.); DEB-2110404 (A.R.); in part by the Great Lakes Bioenergy Research Center, U.S. Department of Energy, Office of Science, Biological and Environmental Research Program under Award Number DESC0018409 (C.T.H.); and the National Institute of Food and Agriculture, United States Department of Agriculture, Hatch project 7005101 (to C.T.H.). C.T.H. is an H. I. Romnes Faculty Fellow, supported by the Office of the Vice Chancellor for Research and Graduate Education with funding from the Wisconsin Alumni Research Foundation. Research in A.R.’s lab is also supported by the National Institutes of Health/National Institute of Allergy and Infectious Diseases (R01 AI153356).

## Supplementary Figures

**Figure S1.**
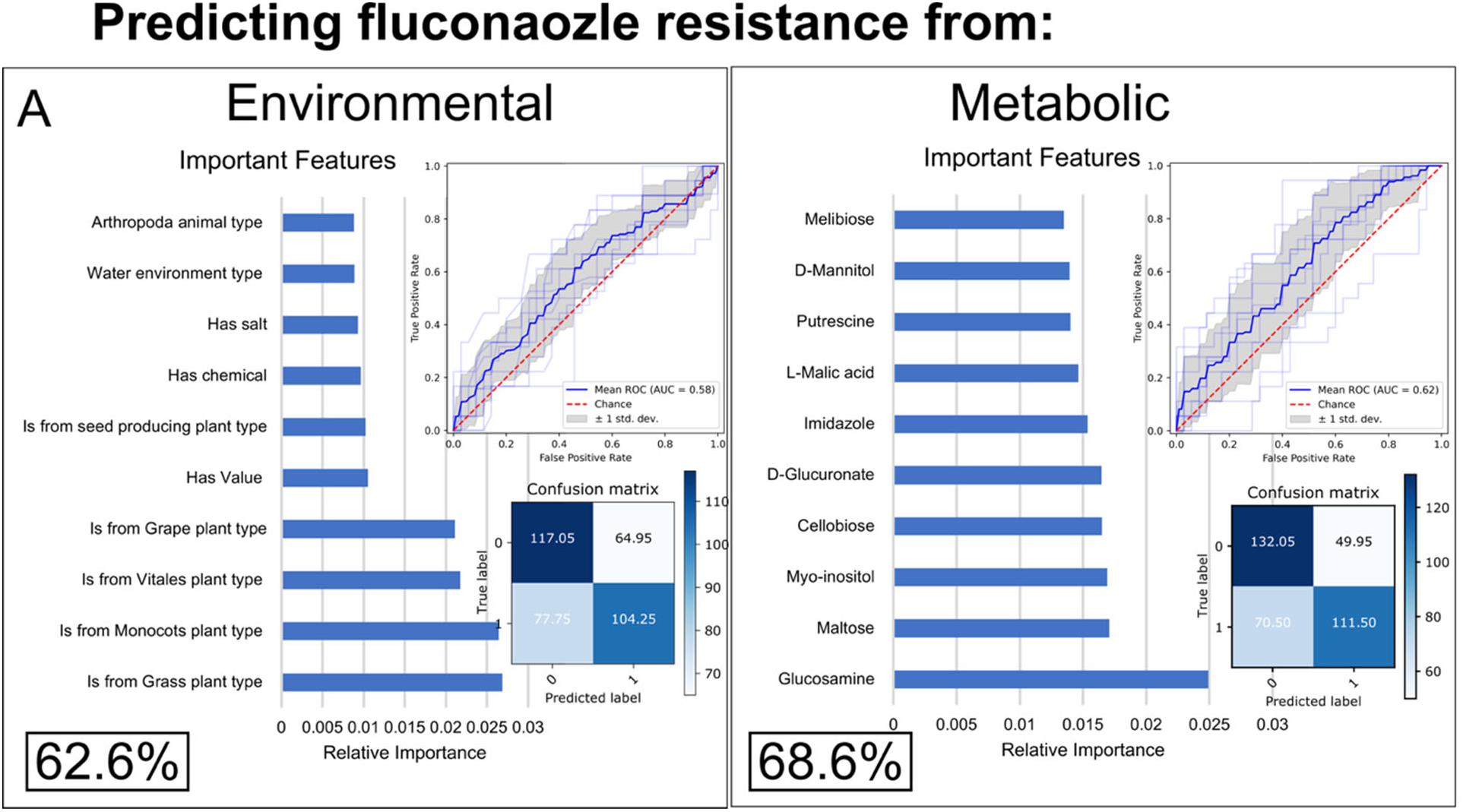
A random forest algorithm weakly predicts fluconazole resistance from environmental and metabolic traits. Accuracy is shown in the form of confusion matrices (bottom right of each panel), which show yeasts predicted correctly to be sensitive to fluconazole (true negatives, top left corner of the matrix), yeasts predicted to be resistant but are not (false positives, top right), yeasts correctly predicted to be resistant (true positives, bottom right), and yeasts correctly predicted to be sensitive (false negatives, bottom left). Receiver Operating Characteristic (ROC) curves (top right of each panel)) show the true positive rate over false positive rate with changing classification thresholds. Feature importance graphs (left of each panel) show the environmental and metabolic features that are most useful for predicting growth on fluconazole. The accuracy in the bottom left corner of each graphic is cross-validated balanced accuracy over 20 down-sampled runs.

**Figure S2.**
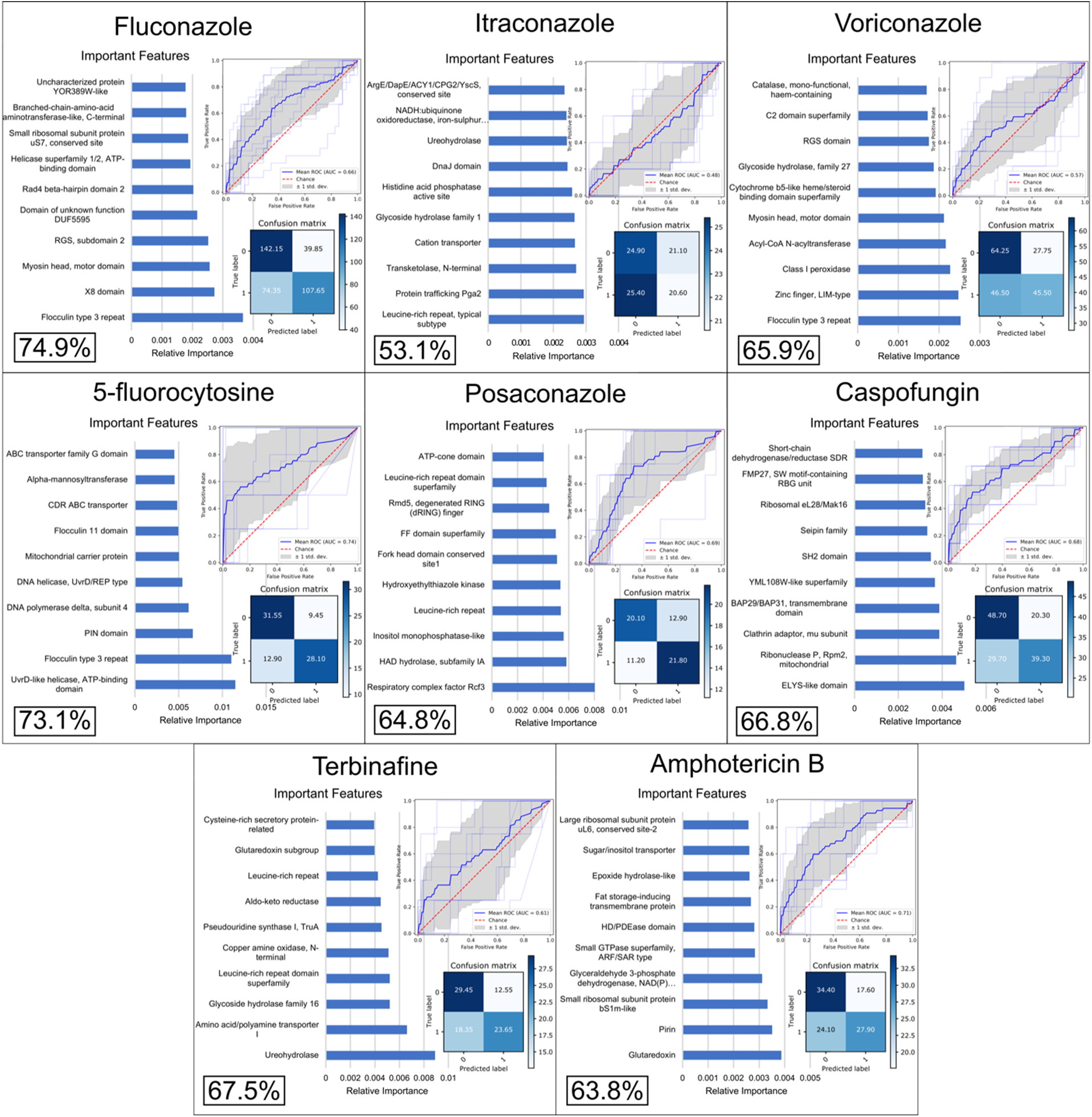
A random forest algorithm predicts resistance to eight antifungal drugs with moderate accuracy from variation in InterPro functional annotations. Accuracy is shown in the form of confusion matrices on the bottom right of each panel, which show yeasts predicted correctly to be sensitive (true negatives, top left of each matrix), yeasts predicted to be resistant but are not (false positives, top right), yeasts correctly predicted to be resistant (true positives, bottom right), and yeasts correctly predicted to be sensitive (false negatives, bottom left). Receiver Operating Characteristic (ROC) curves (top right of each panel) show the true positive rate over false positive rate with changing classification thresholds. The bottom left of each panel corresponds to the average cross-validated balanced accuracy over 20 down-sampled runs. Feature importance graphs (left of each panel) show the InterPro annotations that are most useful for predicting growth on the two drugs. Note that the most informative genomic features were not linked to known drug resistance genes.

**Figure S3.**
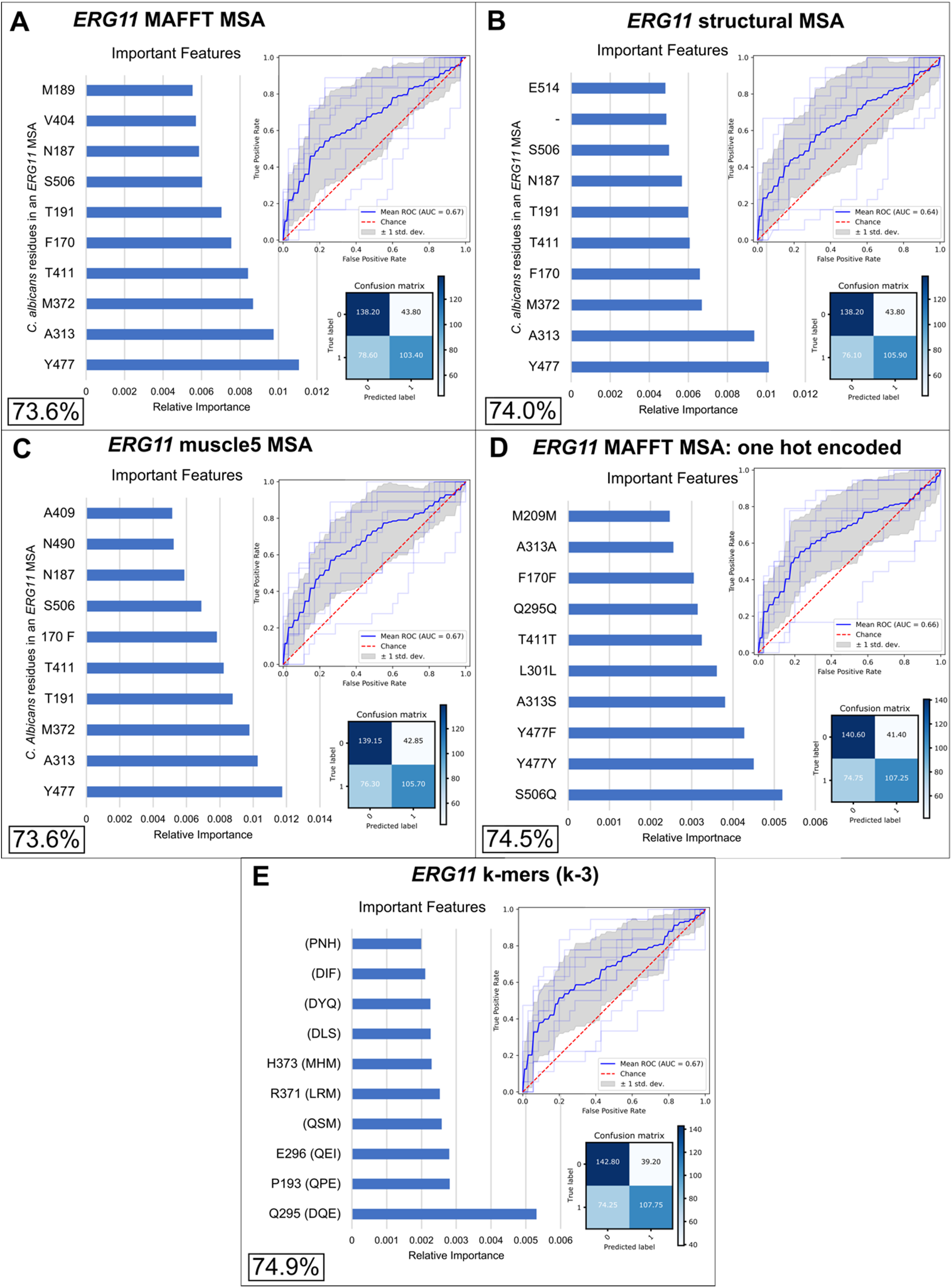
Different Erg11 alignments and ways of encoding sequence information are similarly accurate for predicting fluconazole resistance and highlight the same sites. Accuracy is shown in the form of confusion matrices (bottom right of each panel), which show yeasts predicted correctly to be sensitive to fluconazole (true negatives, top left of each matrix), yeasts predicted to be resistant but are not (false positives, top right), yeasts correctly predicted to be resistant (true positives, bottom right), and yeasts correctly predicted to be sensitive (false negatives, bottom left). Receiver Operating Characteristic (ROC) curves (top right of each panel) show the true positive rate over false positive rate with changing classification thresholds. The accuracy in the bottom left corner of each graphic is cross-validated balanced accuracy over 20 down-sampled runs. Feature importance graphs (left of each panel) show the sites and variants are most useful for predicting resistance to fluconazole.

## Supplementary Tables

**Table S1. Isolation environments of each mammalian-associated yeast in the antifungal drug resistance dataset.**

**Table S2. Resistance to eight different antifungal drugs for 532 species of *Saccharomycotina* yeasts.**

**Table S3. Frequency of resistance to each antifungal drug in each order of *Saccharomycotina*.**

**Table S4. Accuracy of predicting resistance to each antifungal drug when a random forest algorithm is trained on metabolic or environmental datasets.**

**Table S5. Average most informative features when a random forest algorithm is trained on metabolic or environmental datasets to predict resistance to eight different antifungal drugs.**

**Table S6. Gini importance of each site in the MAFFT alignment of all Erg11 protein sequences across *Saccharomycotina* yeasts, which *C. albicans* residue they correspond to (if any), and whether (*) that site has been previously observed in clinical isolates.**

**Table S7. All 36 clinical variants known to conference resistance, the study that identified them, their count numbers in the Y1000+ Project dataset, and their Gini importance for each different method of encoding Erg11.**

